# Monitoring the prolonged TNF stimulation in space and time with topological-functional networks

**DOI:** 10.1101/803817

**Authors:** Stylianos Mavropoulos Papoudas, Nikolaos Papanikolaou, Christoforos Nikolaou

## Abstract

Genes in linear proximity often share regulatory inputs, expression and evolutionary patterns, even in complex eukaryote genomes with extensive intergenic sequences. Gene regulation, on the other hand, is effected through the co-ordinated activation (or suppression) of genes participating in common biological pathways, which are often transcribed from distant loci. Existing approaches for the study of gene expression focus on the functional aspect, taking positional constraints into account only marginally.

In this work we propose a novel concept for the study of gene expression, through the combination of topological and functional information into bipartite networks. Starting from genome-wide expression profiles, we define extended chromosomal regions with consistent patterns of differential gene expression and then associate these domains with enriched functional pathways. By analyzing the resulting networks in terms of size, connectivity and modularity we can draw conclusions on the way genome organization may underlie the gene regulation program.

We implement our approach in a detailed RNASeq profiling of sustained TNF stimulation of mouse synovial fibroblasts. Bipartite network analysis suggests that the cytokine response set by TNF, progresses through two distinct transitions. An early generalization of the inflammatory response, marked by an increase in related functions and high connectivity of corresponding genomic loci, that is followed by a late shutdown of immune functions and the redistribution of expression to developmental and cell adhesion pathways and distinct chromosomal regions.

Our results suggest that the incorporation of topological information may provide additional insights in the underlying topological constraints that are shaping gene expression.

## 1. Introduction

From the analysis of the first genome-wide expression experiments, it became clear that expression levels were related to the organization of genes in linear dimension (1). More recent studies have shown that transcriptional activation may spread in “waves” that affect nearby genes (2) and that a significant proportion of gene expression events may be attributed to genomic position (3). Such tendencies are reflected on evolutionary constraints, with genes in close proximity showing similar patterns of evolution (4–6). These constraints inevitably lead to genes, involved in common functional pathways to be lying closer to each other in the linear genome (7, 8), but also in the genome’s three dimensional structure (9). These preferences are more pronounced in vertebrate mammals and insects (6). They often overlap with characteristic epigenetic modification patterns (10, 11), different combinations of which have been shown to delineate epigenetic “chromatin states” that reflect different levels of regulatory and transcriptional activity (12–14). Overall, there is an increasing interest in the concept of genome compartmentalization focusing on its underlying regulatory and transcriptional activity, evident in recent approaches on cis-regulatory domains (15) and in attempts to model gene expression levels on the basis of genomic position (3). The biomedical importance of this organization is particularly important in various types of cancer, where extensive genomic translocations are the main cause for aberrant regulation of genes and eventually for pathogenesis (16).

Despite the large number of indications for the existence of neighbour effects in the regulation of gene expression, the study of possible underlying mechanisms has been limited. Early works in the simple eukaryotic genome of *S. cerevisiae* have shown the existence of regions of gene expression correlation (17) and referred to their dynamics (18). At the level of functional gene regulation, we have sufficient knowledge of how transcription programs are performed through the activation of specific pathways and gene regulatory networks (19). We also have a variety of tools for assessing the importance of cellular processes and biological pathways through functional analyses of gene expression (20, 21). Nevertheless, there is a growing need for methods that will incorporate spatial information and, until recently, the use of chromosomal linear distance as a predictive marker of gene expression has been limited to a few, large-scale projects with availability of a large number of additional measurements (22).

In the past we have studied the positional footprint on gene deregulation in models of genome compartmentalization in yeast (23, 24), where we have shown both transcriptional regulation and local nucleosomal structure to be delineating distinct genomic domains. In this work we propose an approach that can act complementary to existing functional enrichment analyses (20, 25, 26) with an additional layer of information coming from the level of genome organization. By analyzing the spatial distribution of differential gene expression, we define regions with consistent gene deregulation profiles, that form extensive (of the order of Mbp) chromosomal domains, where limited fluctuation of gene expression changes is suggestive of underlying organizing principles. Association of these regions with functional terms and pathways, through typical gene enrichment analyses, leads to the creation of topological-functional bipartite networks that may then be studied at multiple levels. We showcase this concept in the context of a well-studied gene regulatory program, the activation of cells that is triggered by TNF.

TNF is the archetype of major activating cytokines, which orchestrate the process of the inflammatory response. The succession of steps upon TNF induction has been shown to involve dynamic RNA turnover (27, 28) but to be also accompanied by changes in the chromatin landscape (29) and the organization of transcription factories (30). Sustained expression of TNF can have devastating effects as is evident in transgenic animal models of inflammatory diseases (31). We have recently analyzed the transcriptome dynamics of fibroblasts originating from such an animal model (32) to show that a gradual activation of inflammatory and immune-related pathways accurately mirrors the phenotype of progressive inflammatory polyarthritis. While valuable from the biomedical viewpoint however, *in vivo* approaches do not allow the precise modeling of gene regulation due to cellular cross-talk. In this work, we chose to focus on an *in vitro* gene expression profiling of synovial fibroblasts, a particular cell type that has been shown to be a key receptor of cytokine cues and a very likely effector of inflammatory diseases such as rheumatoid arthritis (31).

Through the analysis of time-course gene expression profiles with the use of positional enrichments and topological/functional bipartite networks, we monitor the progression of the cytokine response in space and time. We show that fibroblasts under prolonged TNF exposure undergo two major transitions that are reflected not only in the activated pathways but also in the clustering of deregulated genes in particular genomic domains.

## 2. Methods

### 2.1 Gene Expression Profiling

Mouse synovial fibroblasts were isolated from C57BL/6 littermate mice. All animals were housed under specific pathogen–free conditions. Three biologic replicates were isolated per experimental condition, and for each condition, a mixed-sex pool of 3 mice was used. Purity of all isolations was assessed by fluorescence-activated cell sorting, with the following acceptance criteria: .85% positive for CD90.2 and, 2.5% positive for CD45. RNA was extracted from mouse synovial fibroblasts with the use of an Absolutely RNA Miniprep kit (Agilent Technologies) and from human synovial fibroblasts with use of a miRNeasy Mini kit (Qiagen). All library preparations, next-generation sequencing, and quality control steps were performed at the McGill University and Genome Quebec Innovation Centre (Montreal, Quebec, Canada). More specifically, for RNA-Seq, TruSeq RNA libraries were prepared and samples were run on an Illumina HiSeq2000 platform using a 100-bp paired-end setup.

### 2.2 Differential Expression Analysis

RNA-seq was performed with three replicates for 5 different timepoints at 1, 3, 6, 24 hours and 7 days after TNF exposure, alongside a 0h control. Mapping was performed with TopHat2 (33) and differential expression was calculated against the 0h control profile with Cufflinks/CuffDiff (34). Differentially expressed genes were defined on the basis of standard thresholds for analysis with |log_2_(FC)|>=1, p-value<=0.05, after adjusting for multiple comparisons.

Functional analysis was performed with the use of gProfileR (26) through its R implementation. Enrichments were studied at the levels of Gene Ontology (GO), KEGG pathways, Transcription Factors (TF) and Human Phenotypes (HP).

Clustering was performed with agglomerative hierarchical clustering using Ward’s minimum variance criterion. Number of clusters was defined based on a simple elbow rule on the within sums of squares values of a k-means clustering approach. Profile similarity calculation was performed through the calculation of euclidean differences in mean cluster differential expression as described in (35).

### 2.3 Creation of Domains of Focal Deregulation

We implemented a method based on unbiased recursive partitioning as described in (36). Differential gene expression data were used as values and their genomic coordinates as a discrete “time-like” variable. A custom R script was written with the used of the R function “breakpoints” from the Package “strucchange” (37, 38). The function performs genome partitioning on the basis of an F-test (Chow Test) applied on consecutive linear models. Once the breakpoints are defined, the script creates a complete partitioning of the genome in discrete regions, each of which is described by a) the number of contained genes and b) their mean differential expression score. An arbitrary criterion of an absolute mean differential expression score >=0.1 was used to call significant DFDs. These were used in the creation of bipartite networks.

Overlaps between DFDs and differentially expressed genes, gene clusters or other sets of genomic coordinates were reported as Jaccard Indexes and assessed statistically through a permutation test performed with a custom script as described in (39).

### 2.4 Topological-Functional Bipartite Networks

These were created in the following way:

1. Starting from a given expression profile, a list of differentially expressed genes is extracted and a set of significant DFDs is called (see above).
2. For each DFD, the differentially expressed genes are being extracted and then passed to gProfileR for gene set enrichment analysis.
3. Functional categories fulfilling significance criteria (Number of Genes in Category >=50, adjusted p-value <=0.05) are associated with the given DFD.
4. The bipartite network is created as an edge list with one vertex being the DFD and the other being its enriched functions.

Networks were analyzed for modularity and visualized with the use of R’s igraph Package (40).

## 3. Results

### 3.1 Complex Patterns of Differentially Expressed Genes in the prolonged stimulation of fibroblasts by TNF

Clustering of the 1595 differentially expressed genes in at least one of the analyzed timepoints revealed some very interesting aspects regarding the prolonged exposure of fibroblasts to TNF. An initial small set (∼330) of deregulated genes (at 1h) is replaced by a much larger (∼640) at 3 hours of stimulation, with less than one third (97) of the genes being shared between the two timepoints. A similar pattern of expression is observed between 3 and 6 hours of stimulation before another abrupt transition at 24 hours with only 55 genes being commonly deregulated between 6h (378 DE genes) and 24h (190 genes). A much longer period extending to 7 days for the last timepoint shows smaller discrepancies in the expression profile. Thus it seems that the state acquired by the cells at 24h remains relatively stable. Overall, the expression data suggest two clear transition points between h1 → h3 (early) and h6 → h24 (late), which may be seen more clearly through the clustering of genes in 8 distinct clusters (Figure 1A, top).

**Figure 1.**
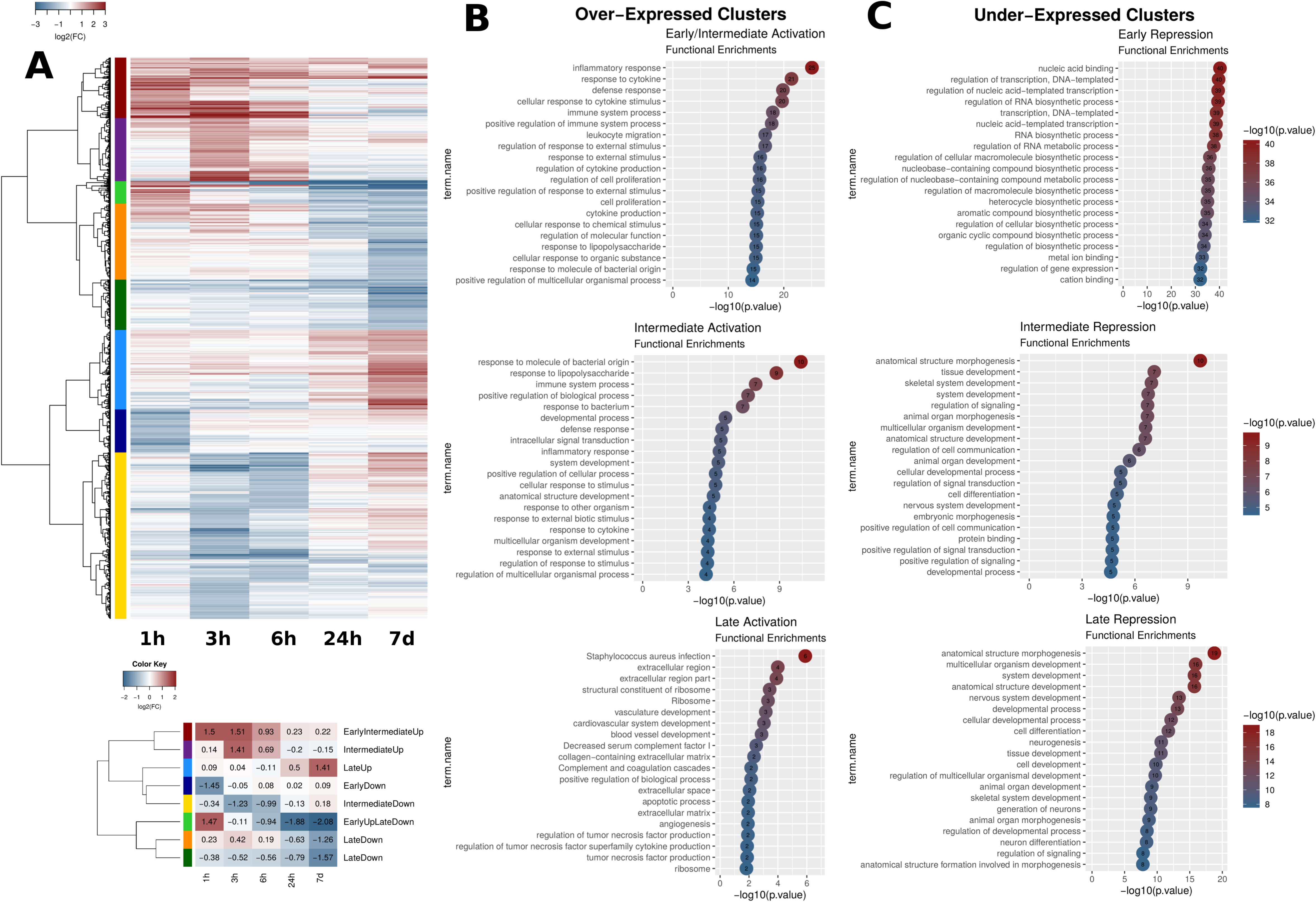
Gene expression clustering and functional analysis. A. (Top) Clustering of gene expression for 1595 genes that were differentially expressed (|log_2_FC|>=1, p.value<=0.05) in at least one timepoint. Red corresponds to over- and blue to under-expression. The 8 clusters created are shown in different colours in the left side of the heatmap. (Bottom) Summarization of the extended heatmap with mean differential expression value for each cluster. Clusters are shown in the same colour coding as above. Cluster names on the right correspond to a general description based on their expression pattern. B, C. Functional analysis of over- (B) and under- (C) expressed clusters. Names of clusters are the same as in (A), with the exception of Late Repression where both “Late Down” clusters from (A) are pooled together. Enriched terms were deduced from a gProfileR analysis. The top 20 enriched terms on the basis of p-values are reported for each cluster. (Functional analysis of the same type for the “EarlyUpLateDown” cluster provided as separate Supplementary Figure 1).

The aforementioned clustering helps us distinguish between genes that are initially over-expressed but gradually restored to normal (unstimulated) levels (Dark red cluster), or even reversed to under-expression (Light green cluster). More importantly, we find most of the clusters to be reflecting clear timepoint-specific expression tendencies. Thus there are clearly under-expressed clusters for early (h1, Dark blue cluster), intermediate (h3 and h6, Yellow cluster) or late (h24, d7, Orange and Dark Green Clusters) timepoints. Similarly the Purple and Light Blue clusters are also reflecting time-specific intermediate (h3 and h6) and late (h24 and primarily d7) over-expression respectively (Figure 1A, bottom).

### 3.2 Functional Enrichment Analysis suggest two points of transition in the cytokine response

Clustering of gene expression profiles allows us to better dissect the dynamics of the implicated functions. We thus analyzed the functional enrichments of the genes in each of the 8 clusters. A clear progression of gene activation may be seen in the functional enrichments of the over-expressed gene clusters (Figure 1B). Going from early (1h) to intermediate (3h, 6h) to late (24h, 7d) timepoints, the functions are shifting from an initial, acute inflammatory response, to more generalized functions of the immune system, to finally become associated with functions related to developmental pathways, apoptosis and the extracellular matrix. It thus appears that fibroblasts, after initially sensing the cytokine cue of TNF, undergo a slow process of switching on major developmental and apoptotic pathways. Given our limited time resolution we can position this transition sometime between 6h and 24h after the initial stimulation.

The shift to developmental functions is also apparent in the down-regulated gene clusters (Figure 1C). Under-expression is strongly associated with transcriptional regulation in the early stage (1h after stimulation). Clusters related to under-expression after 3h are mostly associated with developmental functions, while those that are specific to the later stages are additionally related to cell-cell interactions such as adhesion and migration. This may suggest that an initial regulatory program gets underway since relatively early. It crystalizes, later on, into major transfoming functions that significantly alter the properties of the cells. One gene cluster (Light Green) is particularly interesting in the sense that it reflects this gradual transformation, containing genes that are very over-expressed in the early timepoint, eventually becoming almost completely repressed after 24h. This cluster is strongly enriched in primary metabolic pathways as well as transcriptional regulation (Supplementary Figure 1). The fact that its genes are readily restored to normal levels (at 3h) and then, subsequently suppressed makes the hypothesis of a re-setting of the initial response rather plausible.

### 3.3 Domains of Focal Deregulation (DFD) reflect spatial preferences of gene expression

Functional enrichment analyses are important but they cannot provide insight into the mechanisms, with which the cells utilize their genome in effecting dynamic changes in the regulatory program. We thus set out to investigate the positional aspect of the gene expression process using a topological enrichment approach. Our goal was to identify chromosomal regions with consistent differential expression in our search for links between the genome architecture and the effected gene expression program.

Through a computational approach inspired by signal processing and applied to gene expression data (see Methods) we were able to create segmentation maps of the genome based on the underlying differential gene expression values. An average of ∼300 such regions were defined in each of the five conditions in our dataset, each of which was assigned with a mean differential expression score, directly calculated from the values of the genes it contained. Regions with increased negative or positive score correspond to areas where gene deregulation is topologically consistent. By setting an arbitrary threshold of a mean absolute score of 0.1, we defined the most significantly enriched of these regions, which we will from hereon refer to as Domains of Focal Deregulation (DFD). The DFD chromosomal positions, size in bases as well as their mean expression scores are visualized in Figure 2A.

**Figure 2.**
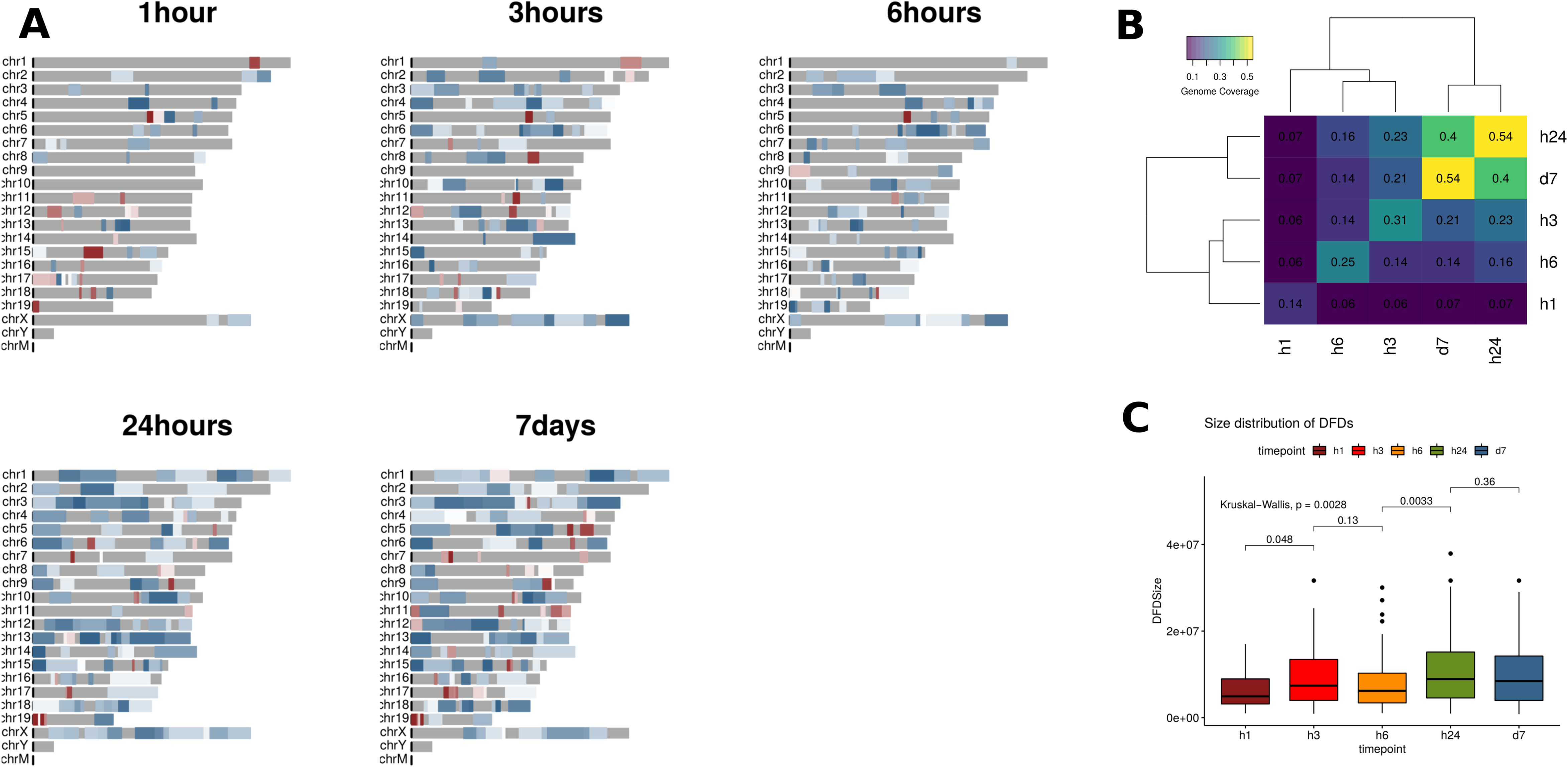
Domains of Focal Deregulation. A. Mouse genome Domainograms showing significant Domains of Focal Deregulation (DFD) with a mean absolute score >= 0.1. Red is positive (over-expression), blue is negative (under-expression) color-coded for intensity. The expansion of the size and number of DFDs is evident, as is the predominance of negative domains, suggestive of an increased clustering of under-expressed genes. B. Genome coverage by DFD as a function of time. Values in the diagonal correspond to genome coverage for each timepoint separately, while the rest of the values correspond to the percentage of the genome covered by DFDs that are overlapping between timepoints. Notice how coverage is clustered in three groups (early:1h, intermediate: 3h, 6h and late: 24h, 7d). C. Size distributions of DFDs for different timepoints. Significant expansions in the size of the DFDs (p.value<=0.05) are observed for the two transition points (1h →3h and 6h →24h).

The Domainograms of Figure 2A reveal significant differences between the five timepoints. Few DFDs in the early timepoint (1h) undergo two waves of expansion at the already discussed transition points of 3h and 24h. At the same time, this quantitative expansion in genome coverage is not coming from the same chromosomal regions, as a number of DFDs emerge and others are depleted between consecutive timepoints. This may be seen in Figure 2B, where we plot the percentage of genome coverage at each timepoint, alongside the corresponding coverage percentages of the sets of overlapping DFDs between them.

It is obvious from Figures 2A and 2B that there are two points of expansion, with the percentage of genome covered almost doubling at 3h and 24h. At the same time, the transitions we have underlined from the functional analyses are also reflected in the similarity between DFD distribution, as may be seen in the dendrogram of Figure 2B. Almost 75% of the DFDs are shared between 24h and 7days and more than half are common between 3h and 6h, but less than one third of the early DFDs are overlapping with those in subsequent timepoints.

The expansion of DFDs, associated with the two transition points is also evident in their size distribution as DFDs at 3h and 24h are significantly longer (Figure 2C). This cannot be directly attributed to the extent of differential gene expression as the corresponding DEG numbers are quite different (643 and 190 for 3h and 24h respectively). Rather, it may be that it is a reflection of a general reorganization of the genome at topological level that occurs through the clustering of genes with particular functional roles. In the following we provide evidence in support of this.

In all, we see that changes in gene regulation associated with the transition from 1h to 3h and from 6h to 24h are also reflected in the topological distribution of genes in linear chromosomal space. At both transition points we observe an expansion in the number and size of DFDs. In the following we turn our attention to the study of this spatial enrichment of gene deregulation, coupled with its functional fingerprint.

### 3.4 DFDs reflect variable clustering of differentially expressed genes and are primarily associated with gene repression

Comparison of the extent and size of DFDs in the domainograms of Figure 2A and the numbers of deregulated genes (see Supplementary Table 1) suggests that they are not directly related. The emergence of DFDs is due to a local clustering of differentially expressed genes rather than to their overall abundance. In order to test this we calculated the global and local enrichments of DFDs in differentially expressed genes in each timepoint as described in Methods (Figure 3A). Even though, general enrichment of DEGs in DFD is expected by definition, the enrichment patterns observed show great variability that reflects the progression of gene expression. We found a decreasing degree in the overall enrichment with time suggesting that the general trend is one towards a spreading out of gene deregulation.

**Figure 3.**
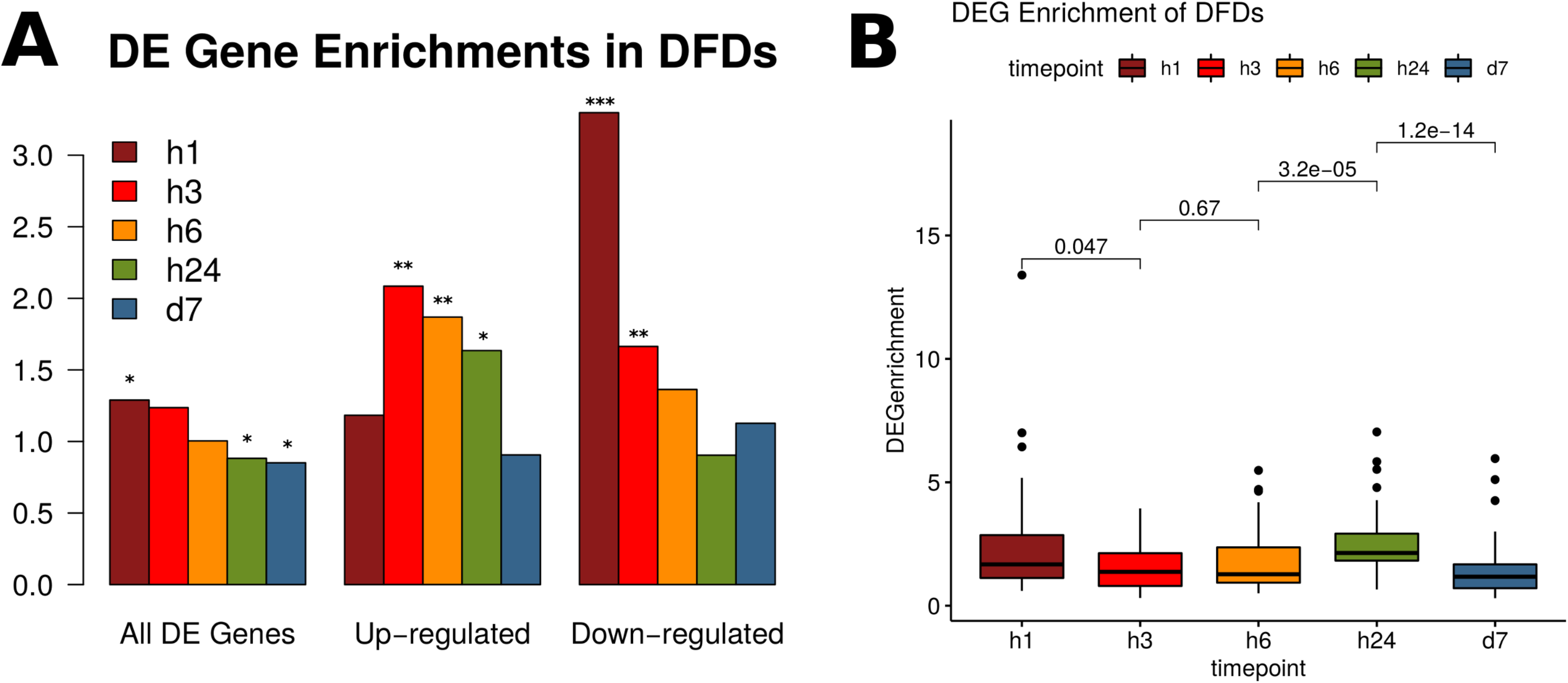
Gene Expression in Domains of Focal Deregulation. A. Overlap enrichments of differentially expressed genes (DEGs) in DFDs for all, over- and under-expressed DEGs. Height of bars corresponds to observed over expected ratios of overlap. Significance levels shown on top of bars are: ***: <0.001, **:<0.01, *:<0.05. B. Distributions of DEG enrichment values on a per DFD basis. Significant drops are observed between 1h → 3h, 6h → 24h and 24h → 7d. The drop in the first transition is towards smaller enrichments (diffusion of DEGs) while the one in the second transition is towards greater enrichments (clustering of DEGs)

Early (1h) DFDs were more enriched in differentially expressed genes but this effect gradually diminished. More interestingly the largest part of DEG clustering occurs in the case of gene repression as may be seen when we plot enrichments separately for over- and under-exressed genes (Figure 3A). Under-expressed genes are highly enriched in the early timepoint, with their preference for DFDs dropping to almost half at the first transition point (3h). In the opposite manner, clustering of over-expressed genes increases in DFDs after the first transition and then also stably returns to the levels of random expectancy. The tendency of under-expressed genes to cluster in DFDs may be also seen directly in the domainograms of Figure 2A (where under-expression is shown in blue and over-expression in red), as well as in the distribution of differential expression scores (Supplementary Figure 3), where the majority of DFDs show significantly low (negative) scores.

The overall enrichments shown above can be masked by effects that are related to the size distributions of DFDs, which as we saw earlier are also significantly variable between timepoints. In order to have a clearer view of DEG clustering, we calculated DEG enrichments at local level, that is for each DFD separately. Here we find a significant clustering at both the initial (1h) and the 24h timepoints (Figure 3B). It thus appears that while the initial timepoint may represent an early bookmarking of particular regions, a general redistribution of differential gene expression occurs at 24h. Notice that in Figure 3B both transitions (1h → 3h) and (6h → 24h) show highly significant changes in terms of enrichments. The first of the two is accompanied by a drop (1h → 3h) while the second by an increase, which are suggestive of different topological tendencies between them. Hence, the transition from 1h to 3h appears to be more one of DFD expansion, with the subsequent “dilution” of DEG enrichment, while the one from 6h → 24h is more likely reflecting a general re-distribution of DFDs with the overal degree of DEG clustering in them increasing.

At a different level, when we analyzed the enrichments of DFDs for genes belonging to the eight time-dependent clusters we also found few but representative enrichments in agreement with the overall tendency for under-expressed genes to cluster in DFDs (Supplementary Figure 4). In this analysis we find early under-expressed genes to be particularly enriched in DFD (see also Figure 3A), while over-expressed genes in both intermediate (3h, 6h) and late timepoints (24h, 7d) show a general avoidance for DFD clustering. These findings are suggestive of different topological clustering tendencies for different functional categories, the study of which is the primary goal of this work. We next turned to the combined analysis of spatial/topological and functional enrichments in gene expression profiles through the introduction of bipartite topological-functional networks.

### 3.5 Topological-Functional Bipartite Networks monitor gene regulation progression

The main hypothesis for this work has been the link between genome organization and the functional outcome of a gene regulation program. Having already established a methodology to assess enrichments at both functional and positional level we went on to combine the two aspects. We defined bipartite positional-functional gene enrichment networks as described in Methods. After constructing bipartite networks for each of the five timepoints, we performed a modularity analysis (41) in order to indentify network modules, which in this context, would represent genomic regions strongly connected with a particular set of functions. Inspection of the resulting networks leads to a number of interesting observations (Figure 4).

**Figure 4.**
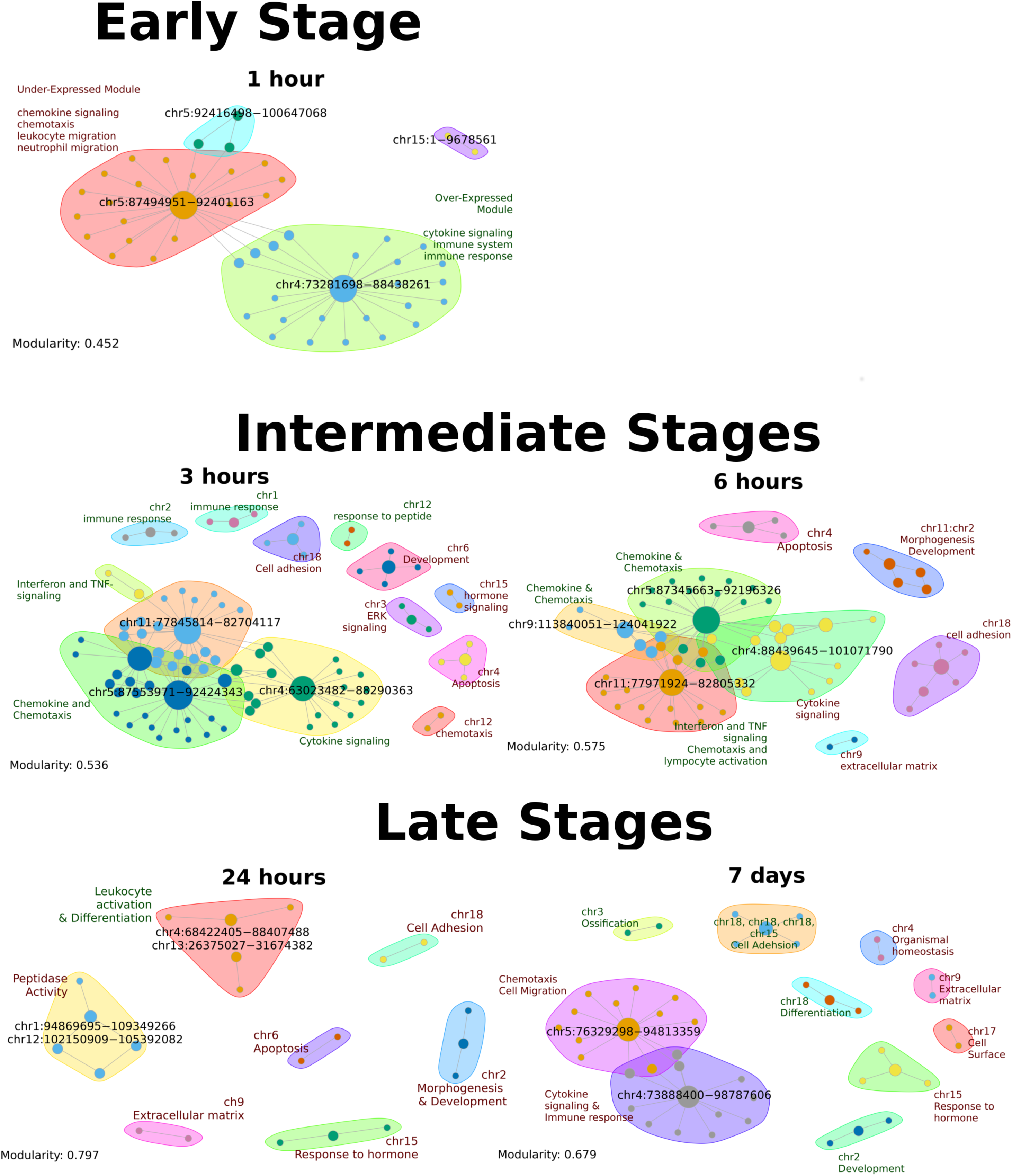
Bipartite Networks for early, intermediate and late stages of stimulation. Bipartite networks for early, intermediate and late stages. Only names of DFD coordinates are shown, while functions are summarised in distinct modules. The expansion with time is evident as is the increase in modularity, which corresponds to a an increasing compartmentalization of functions.

Firstly, that the number of DFDs and associated functions are, to some extent, reflected on the size of the networks. The early (1h) consists of only four modules while the late (24h and 7d) contain 7 and 10 modules respectively. The modularity of the networks is also increasing with time, raising from 0.45 to 0.80 for 24h, which is suggestive of stronger links between chromosomal regions and associated functions as time progresses.

More detailed examination of the networks reveals a number of key elements related to the way cells are affected by the prolonged stimulation by TNF. Initial stimulation (1h) affects a pair of core modules associated with cytokine and chemokine signaling. These are located in chromosomes 4 and 5 respectively but they are not both affected in the same way. The cytokine module (chr4) consists primarily of over-expressed genes, while the chemokine and chemotaxis is enriched in under-expressed genes.

This core pair of modules is also part of the first of the intermediate networks (3h), in which it is expanded through the addition of an interferon/TNF-related module (chr11). One major change in the 1h → 3h transition is the activation of the chemokine/ chemotaxis module, which leads to the formation of a core of three activated modules (cytokine, chemokine and interferon) bridging parts of chromosomes 4, 5 and 11 respectively. A set of additional, smaller modules, associated with the immune response, development and apoptosis also arise at this stage. Progression to 6h brings about two major changes. First, an expansion of the chemokine module with an additional module associated to chr9 and second, the switch of the cytokine module, which still forms part of the network core, to under-expression. This is an indication of the first signs of a shutdown of inflammatory functions which will be generalised in later stages. A set of under-expressed modules related to cell adhesion, apoptosis, development complete the network.

The second major transition from 6h to 24h is accompanied by the complete disappearance of the cytokine-chemokine-interferon core. The highly modular 24h network is the most fragmented one, comprising a set of isolated modules among which a number of functions such as cell adhesion, development and differentiation are associated with under-expression. These last three modules become over-expressed in the network of day 7, in which, we additionally observe the re-emergence of the cytokine-chemokine initial core, now strongly associated with under-expression.

### 3.6 Functional dynamics of bipartite networks are consistent with two transition points at 3h and 24h

Having observed extensive changes in the bipartite networks we went on to assess the changes in a quantitative manner by examining the number and type of functional modules that are emerging, removed, expanded or contracted in the networks. A simple analysis of the number of times a function appears in each network is shown in Figure 5A and is again representative of two major transition points in the system under study. Figure 5A shows an initial, large increase in the number of functions as we move from 1h to 3h. The 3h-acquired functions are related to chemotaxis and interferon signaling as suggested by the network modules. There are overall large similarities in the functional patterns of 3h and 6h, but these are followed by a radical depletion of the largest part of the functions in 24h. Thus, this second transition point is marked by extensive re-organization of functions.

**Figure 5.**
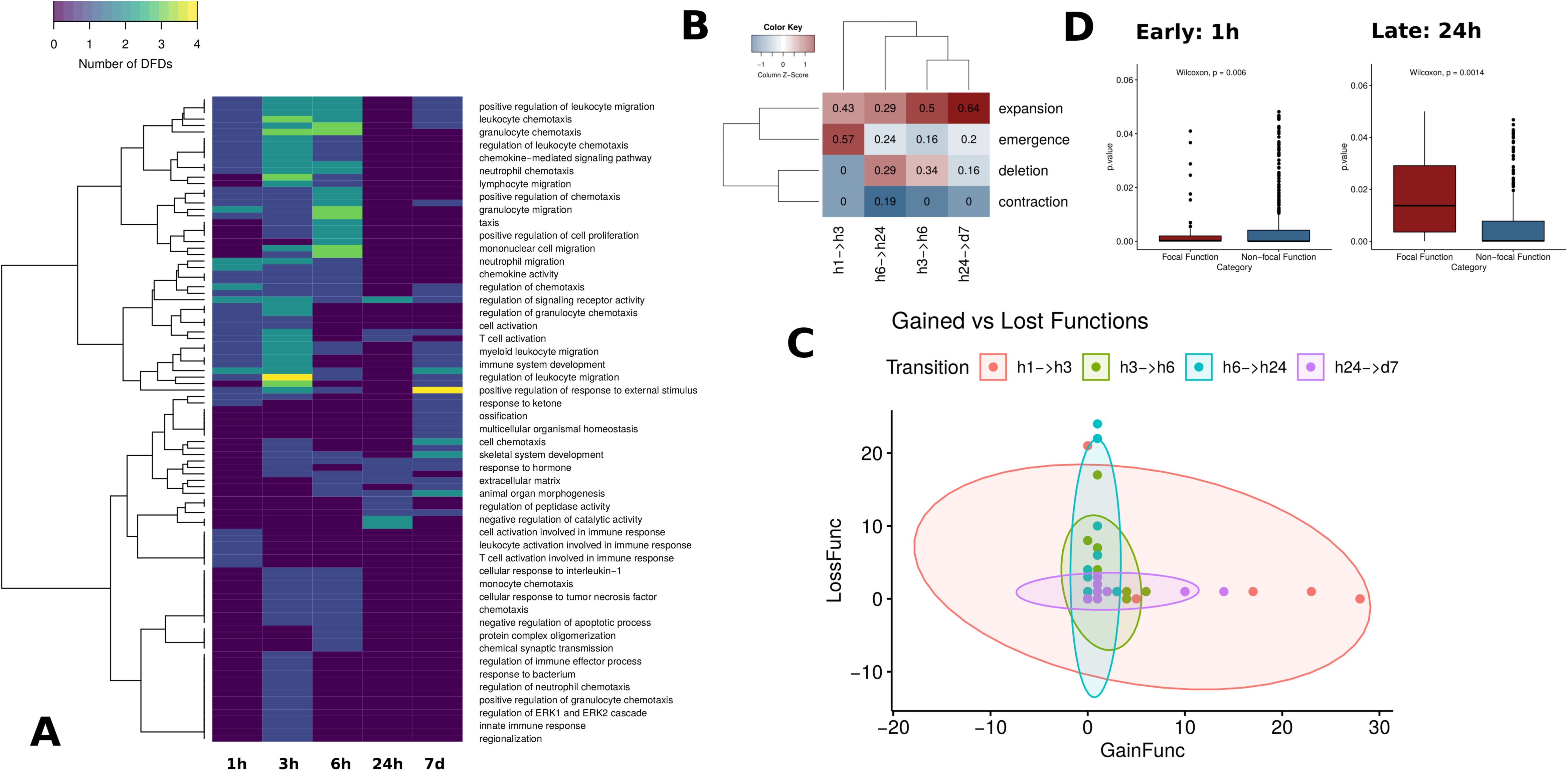
Bipartite network dynamics. A. DFDs numbers per function in bipartite networks. Each cell in the heatmap shows the number of DFDs associated with each function. Functions with 0 DFD (blue) are absent from the corresponding bipartite network. An expansion at 3h and a subsequent decrease and redistribution at 24h is evident. B. Percentage of DFDs belonging to each dynamic category for comparisons between consecutive timepoints. Emergence is prominent for the early transition, while deletion is quite significant for the late one. C. Number of gained/lost functions per DFD in the bipartite network comparisons between consecutive timepoints. h1 → h3 transition is marked by an acquisition of functions while h6 → h24 by a general loss. D. Functions clustered to DFDs behave differently between timepoints. Enriched functions were divided into those associated with a DFD in bipartite networks and those that were not. Mean enrichment of functions that are associated with DFDs is smaller than the one of functions that are not attached to a DFD for the early stage. The situation is inversed for the 24h timepoint.

Most of the inflammation and immune response-related functions are not present at 24h. Instead they have been substituted by pathways associated with development, cell adhesion and cell motility, which are suggestive of major transformations occurring in fibroblasts under prolonged TNF stimulation. A subsequent expansion of similar developmentally-related functions occurs in the latest timepoint (7 days). Most of the intermediate stage (3h, 6h) functions are absent but functional terms such as cell adhesion, cell differentiation and and cell migration are prominent.

### 3.7 Dual dynamics of positional and functional enrichments suggest intermediate expansion followed by late re-organization

The changes in the functional footprint through time, discussed above, are also accompanied with changes in the domains of focal deregulation. As domain boundaries are more difficult to identify directly, we implemented a computational approach based on chromosomal coordinate overlap to assess the qualitative nature of the change. In this way, domains that are present in a given condition (timepoint) but not overlapping any corresponding domain in a subsequent timepoint are considered to be “depleted”, while the opposite are assigned as “emergent”. More complex cases of domains which overlap were identified as either “contracted” or “expanded” depending on whether their boundaries were stretched or withdrawn between consecutive timepoints.

The results of this analysis are shown in Figure 5B, where the percentage of each of the four categories of domain changes was calculated over the total extent of DFD coverage. The two transition points (h1 → h3) and (h6 → h24) show the greatest degree of domain changes but in different ways. The early transition point (h1 → h3) is marked by the emergence of a large number of new domains (>50% of the total), while the rest correspond to expansions. One the other hand, the h6 → h24 transition is the only such that contains contracted domains with a similar proportion of expansion, deletion and emergence signifying a general domain re-organization.

### 3.8 Prolonged TNF stimulation goes through an initial expansion and a late-stage contraction of activated functions

Figure 5A suggests two major shifts in the number of enriched functions between timepoints. Since these changes are tightly linked to DFD dynamics we wanted to test this further by looking into each DFD separately. We calculated the number of functions attributed to each DFD and compared them between consecutive timepoints, distinguishing between “gained” and “lost” functions. We then plotted the corresponding numbers for each DFD and grouping for the four transitions (h1 → h3, h3 → h6, h6 → h24 and h24 → d7) in Figure 5C. The plot shows two distinct trends in the shape of the bubbles (with horizontal and vertical orientations corresponding to predominant gain and loss of functions respectively). One can see that h1 → h3 and h24 → d7 transitions are associated with function gain, while h3 → h6 and h6 → h24 with function loss, while it is also clear that the effects are stronger for the h1 → h3 expansion and the h6 → h24 contraction. Indeed, it is these two transitions that are also statistically significant in terms of number of acquired/lost functions per DFD (Supplementary Figure 5).

An overview of the analyses of our bipartite network approach strongly suggests that the two major transition points in the prolonged stimulation of fibroblasts by TNF are also qualitatively different, with the an early expansion that is probably reflecting the generalization of the immune signaling response, followed by a major shutdown of immune-related pathways in 24h.

### 3.9 Focal functions attract a greater number of differentially expressed genes

Our data suggest complex dynamics of enriched functions, being lost and gained from equally volatile DFDs. This dynamics is, however, confined to the subset of functions that are associated with DFDs. A large number of functional categories are also enriched in differentially expressed genes without being linked to focal deregulation. These are being enriched in genes that are distributed more broadly in genome space and may thus be subject to different expression biases. We tested these differences through a comparison of enrichment p-values for functional terms associated with DFDs (focal functions) against non-focal functions, that are enriched but not linked to particular chromosomal regions (Figure 5D). Interestingly, we find significant differences between early (1h) and late (24h) stages. At the early stage, focal functions are significantly more enriched than non-focal ones, while the opposite holds for the 24h timepoint. This result, also implied by the bipartite network modularity analysis, is suggestive of an increased fragmentation of differential gene expression into distinct regions and functions.

## 4. Discussion

Our work constitutes one of the first attempts to incorporate positional information in the analyses of gene expression. The starting hypothesis is that differential gene expression may be clustered in confined regions of the linear genome due to regulatory, epigenetic and structural constraints. Indeed, elements of the three-dimensional genome structure such as TADs (42) have been shown to delineate genomic space with particular transcriptional tendencies, while we (23) and others (43) have demonstrated focally increased transcription in TAD boundaries. More recent works have focused on the regulatory potential of self-contained linear genomic regions, thus called cis-regulatory domains (CRDs) (15). Through a relatively simple approach, we herein demonstrate that regions of consistent differential expression are common and may, moreover provide insight in the way a gene expression program develops in time.

When addressing the level of differential gene expression clustering in our defined DFDs we find a significant enrichment for under-expressed compared to over-expressed genes. Even though we cannot exclude that it may be a particular characteristic of the system under study, it is worth noticing that repressive domains are readily formed in eukaryotic genomes mediated by Polycomp-Group (PcG) proteins (44) or through the association of genomic regions with the nuclear lamina (45). Thus it will not be surprising to find that there is a stronger overall clustering tendency for repressive genes in order to maintain transcriptional silencing.

Another interesting aspect that comes out of our time-dependent study is related to the progressive expansion of the genome space that is covered by DFDs. Since this is not correlated with the number of differentially expressed genes (coverage by DFDs peaks at 24h, where we have the smallest number of DEGs) we may assume that it reflects a propensity for increased genome compartmentalization. Indeed, we find that, with time and independently of the number of differentially expressed genes, expression tends to be more “focal” and this is, moreover, accompanied by an increase of the bipartite network modularity. Again, while this may be a singular property of the cytokine response, it deserves to be studied in more detail and in different systems.

Besides providing insight on the functional modularity of gene expression profiles, the bipartite networks that we describe in this work may also assist in the formulation of hypotheses on genome organization. Interacting modules observed in the bipartite networks of the intermediate stages (3h and 6h) show strong functional interactions between regions from different chromosomes (chr4, chr5, chr9 and chr11 in particular). It would be really interesting to investigate whether such interactions are also reflected upon the three-dimensional organization of the genome. Even though trans-chromosomal interactions are inherently difficult to detect, new methodological approaches such as SPRITE (46) and GAM (47) would probably allow us to test similar hypotheses.

Another promising aspect of our work is related to the analysis of genes belonging to focal vs non-focal functional categories. Functional categories that tend to have their genes clustered in close proximity are more likely to be enriched depending on the stage of the process under study and this may be an indication of a more focal or more widespread expression program. One interesting question would be to examine genes, whose expression may be attributed to their relative position rather than their participation in a certain pathway. We would call these “by-stander genes” as, in essence, one could suggest that their mis-expression is driven by nearest neighbor effects. Modeling the likelihood for positional vs functional drive of such by-standers is a very interesting prospect for follow up studies.

Overall, our combined analyses of DFDs and bipartite network dynamics strongly support the notion of genome architecture having a fundamental role in the development of gene regulation. Implementation of this approach in different systems and more comprehensive datasets is bound to provide additional insights on the underlying mechanisms.

## Supplementary Material

Supplementary material can be found in the online version of this article.

## Data Availability

Processed Gene Expression data, differential gene expression lists, details on the gene clusters, DFDs and bipartite networks as well as custom R scripts used in the analyses are deposited in Mendeley Data: doi:10.17632/fytpjj5ny5.1.

## Acknowledgments

The authors are grateful to Prof. George Kollias for providing access to experimental facilities at BSRC “Alexander Fleming”, Greece. We would also like to thank Dr. Vangelis Ntougkos for assistance in the performance of experiments and Eleni Lianoudaki for proofreading this manuscript.

## Supplementary Material

**Supplementary Table 1.** Numbers of differentially expressed genes (over-expressed, under-expressed and general total) for each timepoint.

|  | Over-Expressed Genes | Under-Expressed Genes | Total DE Genes |
| --- | --- | --- | --- |
| 1h | 192 | 146 | 338 |
| 3h | 303 | 347 | 650 |
| 6h | 117 | 266 | 383 |
| 24h | 40 | 152 | 192 |
| 7d | 281 | 394 | 675 |

**Supplementary Figure 1.**
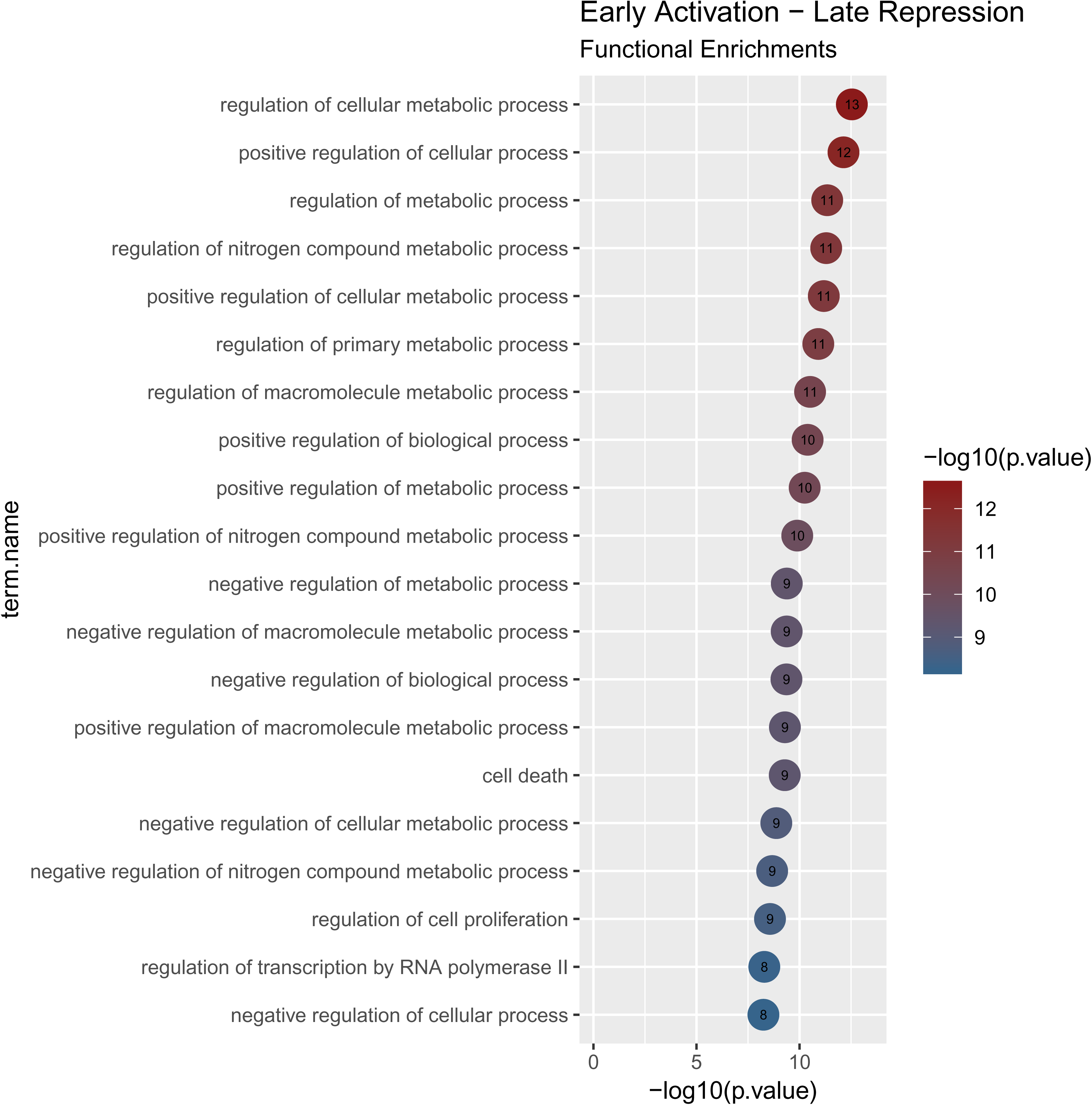
Functional Enrichment analysis of the High-Low expression cluster (top 20 enriched terms). Functionally enriched terms almost entirely belonged to transcriptional regulatory factors affecting primary functions such as metabolism, in particular of macromolecules.

**Supplementary Figure 2.**
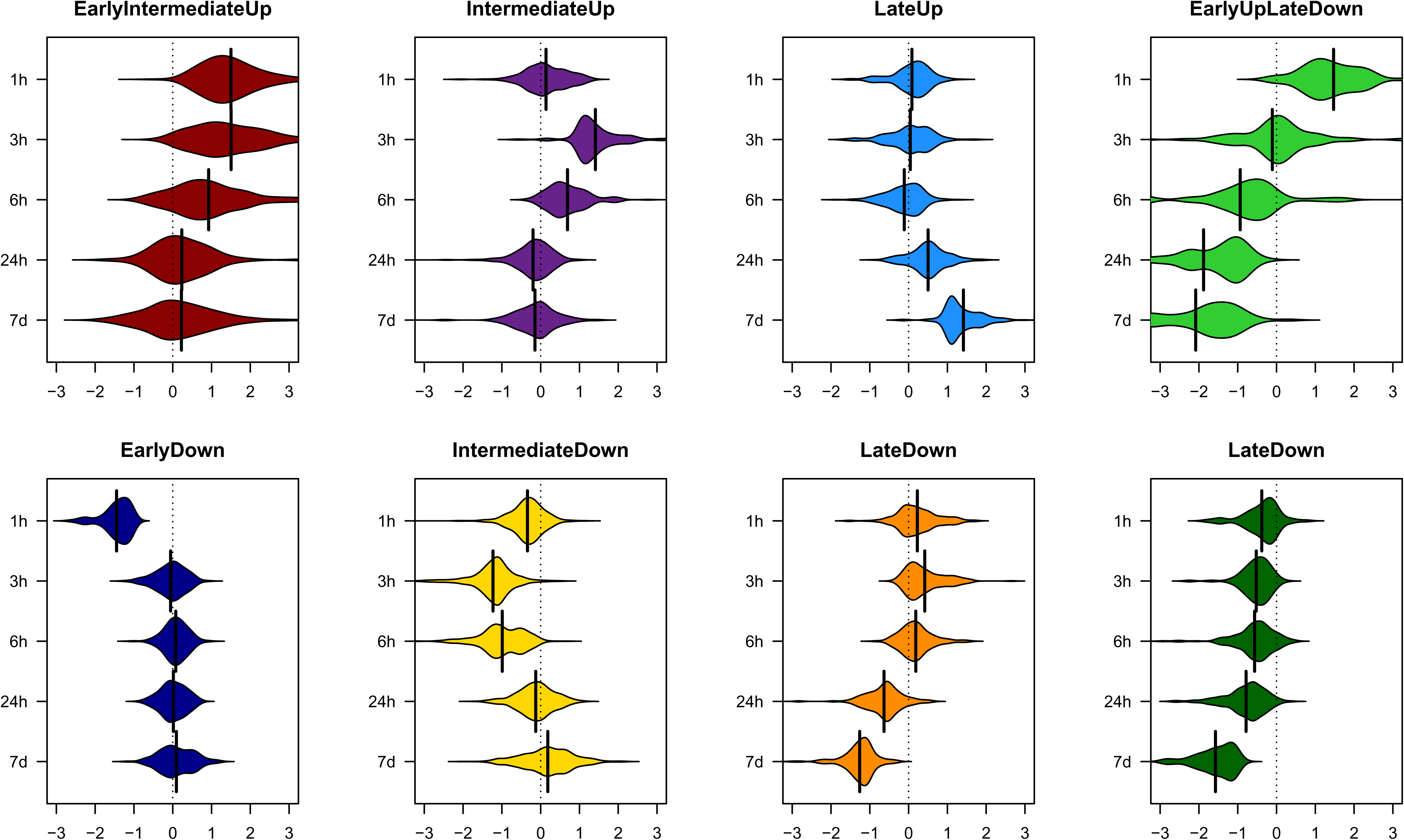
Distributions of differential expression values for each of the 8 defined clusters across timepoints.

**Supplementary Figure 3.**
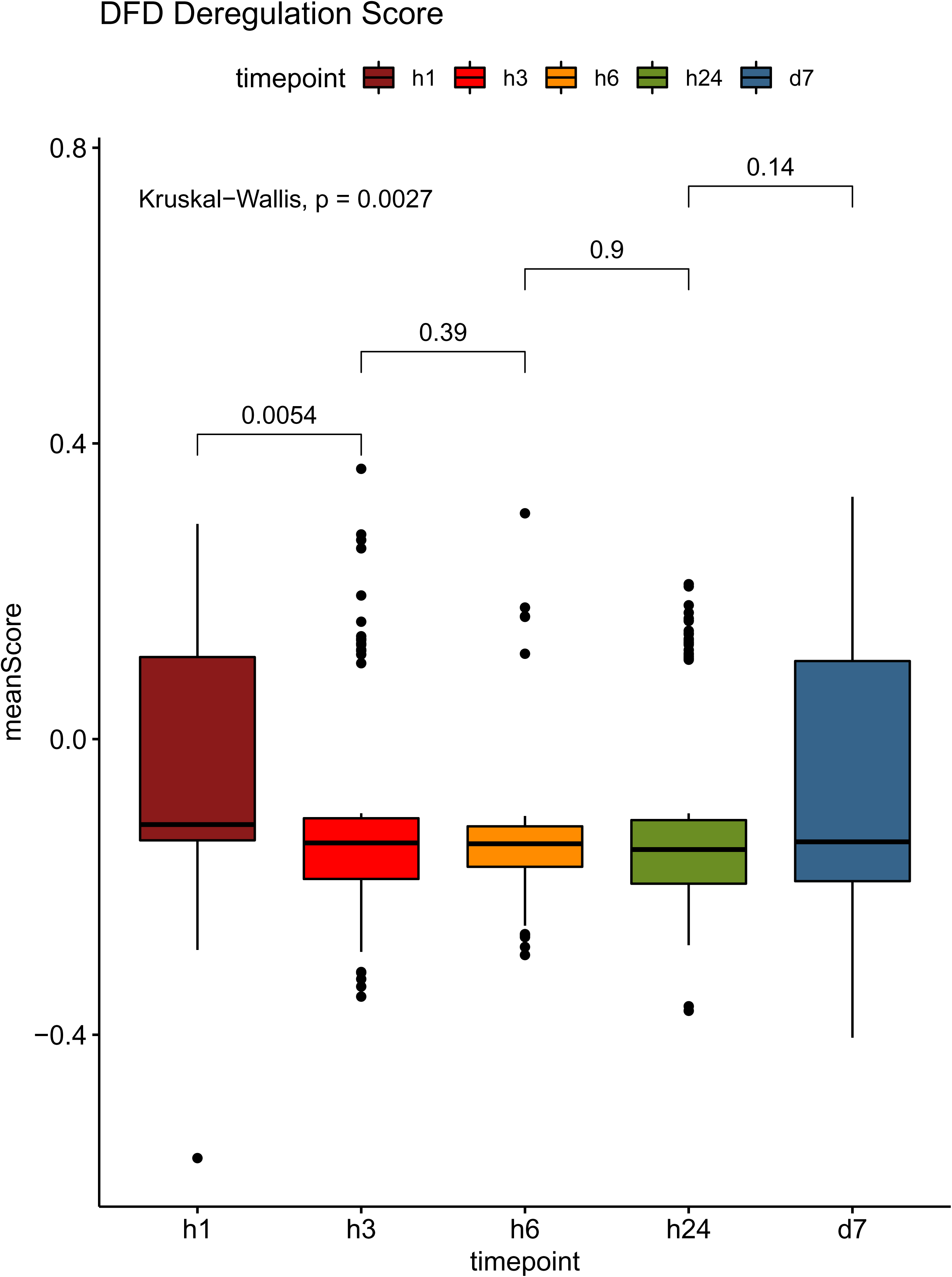
Distributions of mean DFD diffential expression scores across timepoints.

**Supplementary Figure 4.**
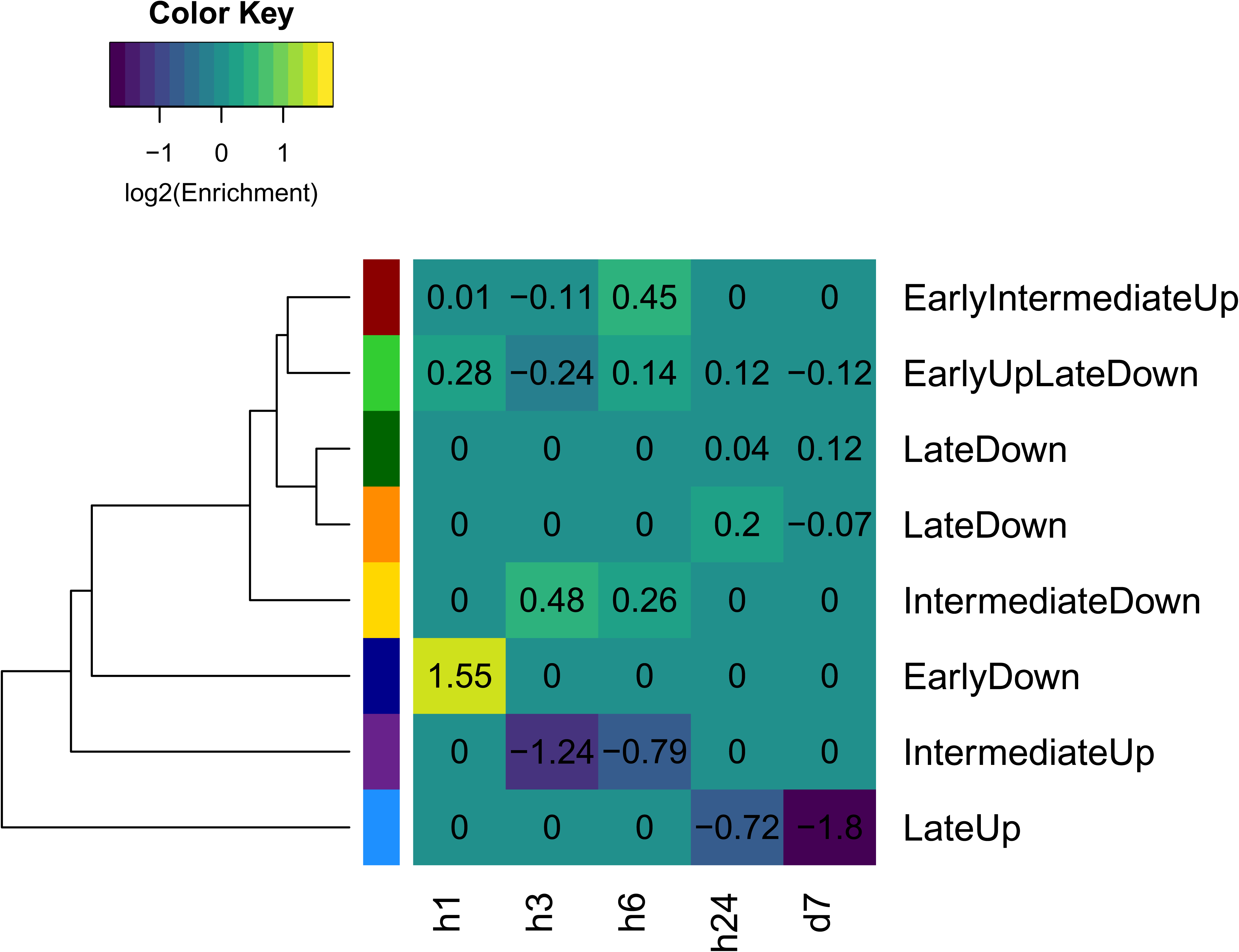
DFD enrichments (in log_2_ scale) for genes belonging in each of the 8 defined clusters. Few positive enrichments are all associated with down-regulated clusters. Negative enrichments, suggesting overall avoidance are associated with up-regulated clusters.

**Supplementary Figure 5.**
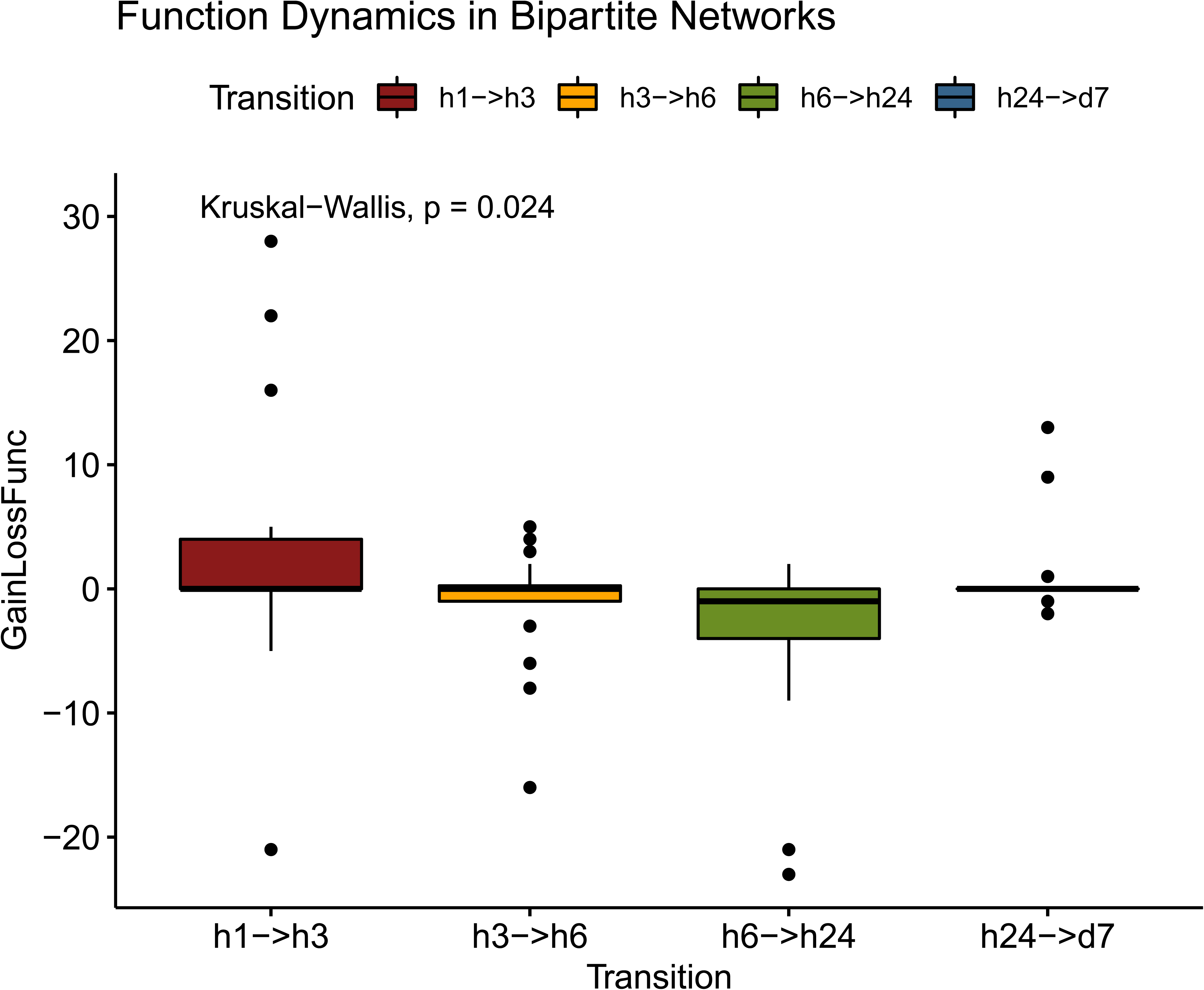
Distribution of the difference between Gained and Lost functions (N_gain_-N_loss_) for bipartite networks of consecutive timepoints. An initial gain for the h1 → h3 transition is reversed to a predominance of lost functions for h6 → h24.

